# An online GPCR drug discovery resource

**DOI:** 10.1101/2025.01.11.632537

**Authors:** Jimmy Caroli, Søren N. Andreassen, Javier Sánchez Lorente, Binghan Xiao, Gáspár Pándy-Szekeres, David E. Gloriam

**Author notes:** Note to reviewers: The resource described in this paper is available at www.test.gpcrdb.com and will be moved to www.gpcrdb.com upon publication.

## Abstract

G protein-coupled receptors (GPCRs) have been targeted across all therapeutic areas, mediate the actions of 516 (36% of all) approved drugs and are being targeted by 337 agents in clinical trials. So far, 121 GPCRs are targets of approved drugs and 30 additional receptors have entered clinical trials and may expand the drugged GPCRome in the coming years. Here, we describe an online resource of GPCR drugs, clinical trial agents, targets and disease indications. This resource offers unique reference data, analysis and visualization, and is availed as a new section, ‘Drugs and Agents in trial’ integrated in the GPCR database, GPCRdb. Furthermore, it includes a target selection tool for prioritization of receptors for future drug discovery. This up-to-date knowledgebase will help identify strategies and trends in current GPCR drug discovery and give insights into which already drugged, or yet untapped targets have the largest potential in specific diseases.

## Introduction

G protein-coupled receptors (GPCRs) form a superfamily of approximately 800 membrane proteins^1^ activated by a diverse array of extracellular stimuli (e.g., odorants, photons, tastants) and endogenous ions, small molecules, peptides, and proteins^2^. Hence, they govern signal transduction in a myriad of physiological and pathophysiological processes, such as sensory perception, hormonal homeostasis, neurotransmission and immune response. Their accessibility at the cell surface, druggable binding sites and abundant modulation of physiology have made GPCRs highly attractive therapeutic targets across all major therapeutic areas. 36% of all approved drugs act on GPCRs^3^, including several drugs listed as essential medicines by WHO (www.who.int/publications/i/item/WHO-MHP-HPS-EML-2023.02) or reaching blockbuster sales, primarily within diabetes and obesity (https://www.drugdiscoverytrends.com/best-selling-pharmaceuticals-2023/).

Recent years have brought several biological and technological advancements that now open new opportunities for the discovery of new drugs targeting GPCRs. Mechanistically, allosteric modulation is strongly increasing in clinical trials^3^ and pathway-biased signaling brings additional potential to increase safety and efficacy, although still awaiting its major translation into more rationally designed clinical agents^4^. Biosensors have been developed with increased sensitivity, throughput and ability to dissect GPCR conformations^5^ and transducers (16 G proteins grouped into four families and four arrestins divided into two β-arrestins and two visual arrestins)^6,7^. Breakthroughs in crystallography and cryo-EM have led to a surge of high-resolution structures of GPCRs (https://gpcrdb.org/structure/statistics) giving valuable insights into ligand recognition^8,9^ and receptor activation^10^. Furthermore, virtual screening scaling to ultra-large molecule libraries have accelerated the identification of novel ligands with favorable potency, target specificity and pathway-biased signaling^11-14^.

These abovementioned advances warrant an updated and comprehensive data repository collating clinical investigations of GPCRs and enabling trends and strategies to be analyzed. Here, we present an open access online resource for compounds, targets and disease indications that are being explored in clinical trials or have reached regulatory approval. This replaces our previous resource from 2017^15^, cover a larger number of data sources and regulatory agencies and has grown to a complete section “Drugs and Agents in Trial” of GPCRdb^16^. Furthermore, it includes a target selection tool aiding the prioritization of GPCRs for drug discovery projects. We believe that these data and tool resources will support drug discovery for both drugged and yet untargeted receptors in specific diseases towards harnessing more of the therapeutic potential of GPCRs and accelerating the development of innovative treatments.

Additionally, this data resource forms the basis of an accompanying review in which we analyze drug discovery trends for the GPCR superfamily^3^. The analysis indicates a continuous rise in the development and approval of GPCR-targeted drugs, targeting both well-characterized and orphan GPCRs, expanding the scope of GPCR-related therapeutics. GPCRs agents are increasingly allosteric modulators and biologics and tested within especially metabolic diseases, oncology and immunology.

## Results

### Overview of the online platform

The online platform can be found in a dedicated section “Drugs and agents in trial” of GPCRdb. It contains three subsections, Agents/drugs, Targets and Diseases, respectively with in total eight pages presenting distinct data and tools (Fig. 1). All these pages are new and replace our resource from 2017^15^, which has been removed. In terms of data, it contains 516 approved drugs, 337 agents in clinical trials, 121 targets of approved drugs and 133 GPCRs, whereof 30 novel targets, in clinical trials. The disease data span 683 disease indications grouped into 23 areas using the International Classification of Diseases (ICD11)^17^.

**Figure 1.**
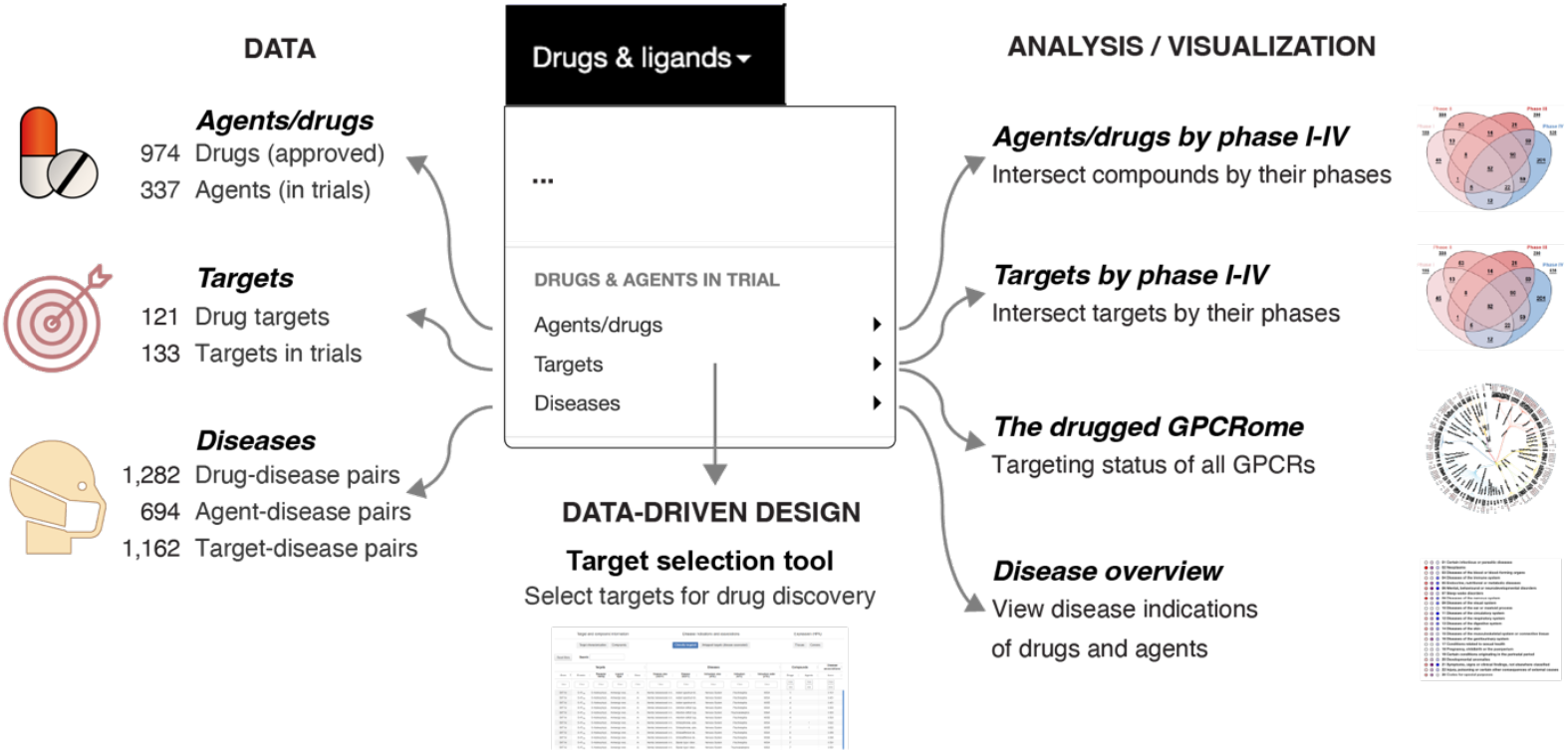
Resource overview. The GPCR drug discovery resource provides reference data and analysis tools for compounds (drugs and agents in trials), targets and diseases across phases I-IV. These data can drive the selection of targets for new drug discovery efforts via a dedicated tool (Fig. 5). The counted target-disease associations (1,162) all have scores >0.5 (data from Open Targets).

### Drugs and agents in clinical trials

#### Agent/drug page

This page provides a full description of an approved drug or agent in clinical trials, which can be retrieved by its name or database identifier (PubChem, ChEMBL, DrugBank and DrugCentral). The top of the page shows the 2D chemical structure, molecule type (e.g., small molecule or peptide), number of indications being tested in each of phase I-III or approvals in phase IV, database links, experimental structure complexes with GPCRs and binding site mutations (Fig. 2). Next, the connections of the compound to different receptor targets and disease indications are visualized in a Sankey diagram. A complete and filterable tabulation of all the compound’s target–disease pairs is shown at the bottom. This starts with information about the activity at different clinical phases (max phase and individual phases) followed by the target. The disease information spans indication names, ICD11 codes, ATC codes and association scores (from Open Targets).

**Figure 2.**
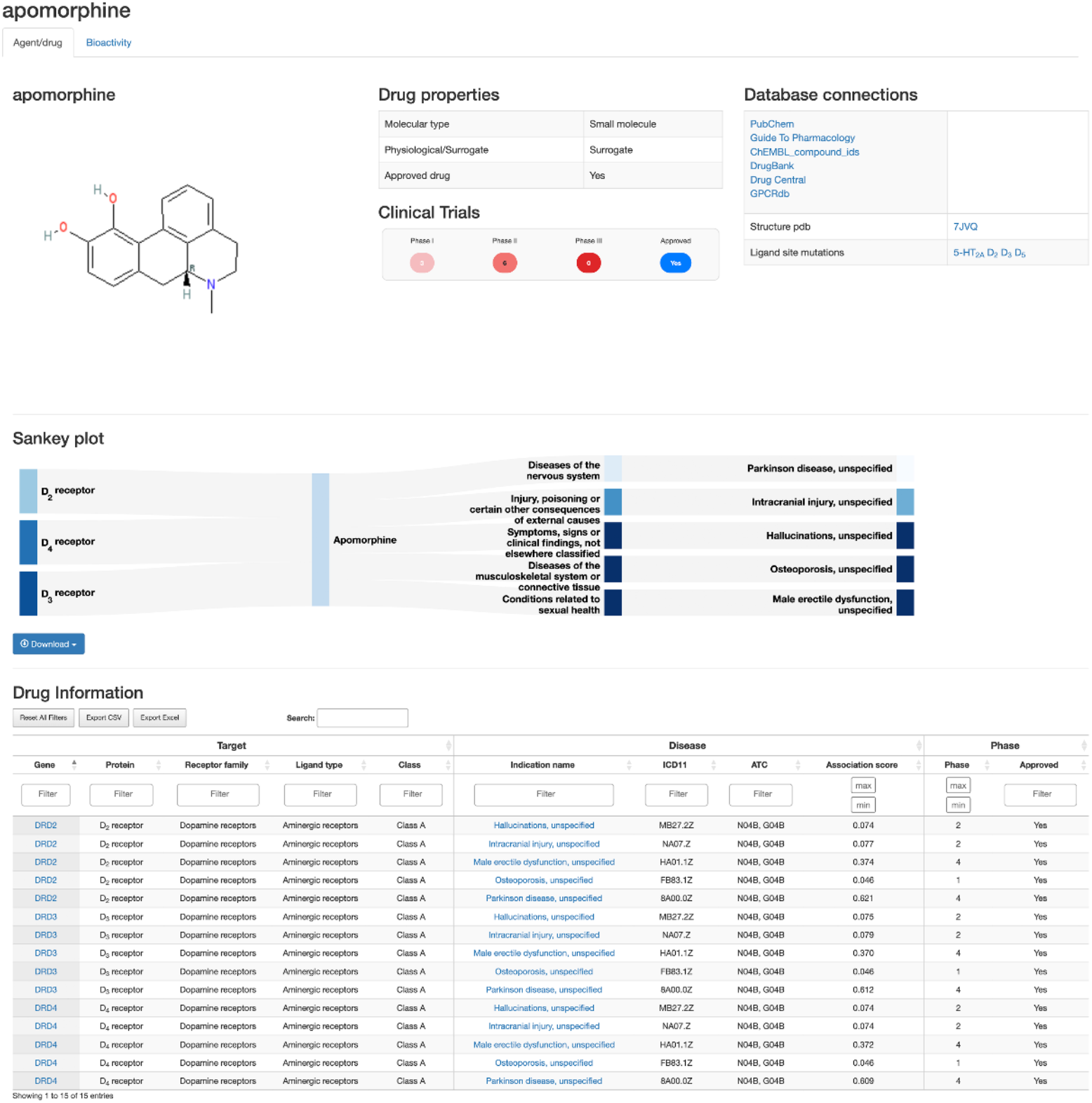
Agent/drug page. The agent/drug page for the approved drug apomorphine exemplifies the different data on chemical structure, molecular properties, database connections, structures, ligand site mutations, targets, diseases and bioactivities (tab not shown here).

A second tab instead shows bioactivities stored in GPCRdb’s ligand database^18^, which aggregates ligands from ChEMBL^19^, Guide to Pharmacology^2^ and the PDSP K_i_ database (https://pdsp.unc.edu/databases/kidb.php). The bioactivity data includes a ligand functional or binding parameter that is presented as minimum, average and maximum values, and the fold selectivity, across all stored experiments. Furthermore, it provides vendors for purchasing and chemical information, including the structure in SMILES notation. While this is here the secondary tab, it is the primary tab instead accessed via the alternative page *Ligands (ChEMBL, GtP, Ki db)* in GPCRdb. The advantage of this is that it allows swift switching between drug-related and bioactivity data, irrespective of the point of entry. Furthermore, it will ensure awareness of additional relevant information, when available (the Agent/drug or Bioactivity tabs is hidden if there is only the other type of data). Altogether, this provides a one-stop-shop for agent and drug information spanning all the relevant information across the compound, target and disease spaces collated from major databases.

#### Agents/drugs by phase I-IV

This page allows compounds to be intersected by one or more clinical phases via a Venn diagram (Fig. 3). Clicking areas of the Venn diagram dynamically updates a table of compounds containing their name, molecule type, pharmacological modality. The target information includes the gene and protein names, as well as classification by class, ligand type and receptor family (by physiological ligand). The disease information include areas and diseases classified using ICD11 codes and ATC codes (drugs only). Hence, whereas the starting point is one or more clinical phases, the compounds of interest can quickly be narrowed down based on the relevant target and disease information.

**Figure 3.**
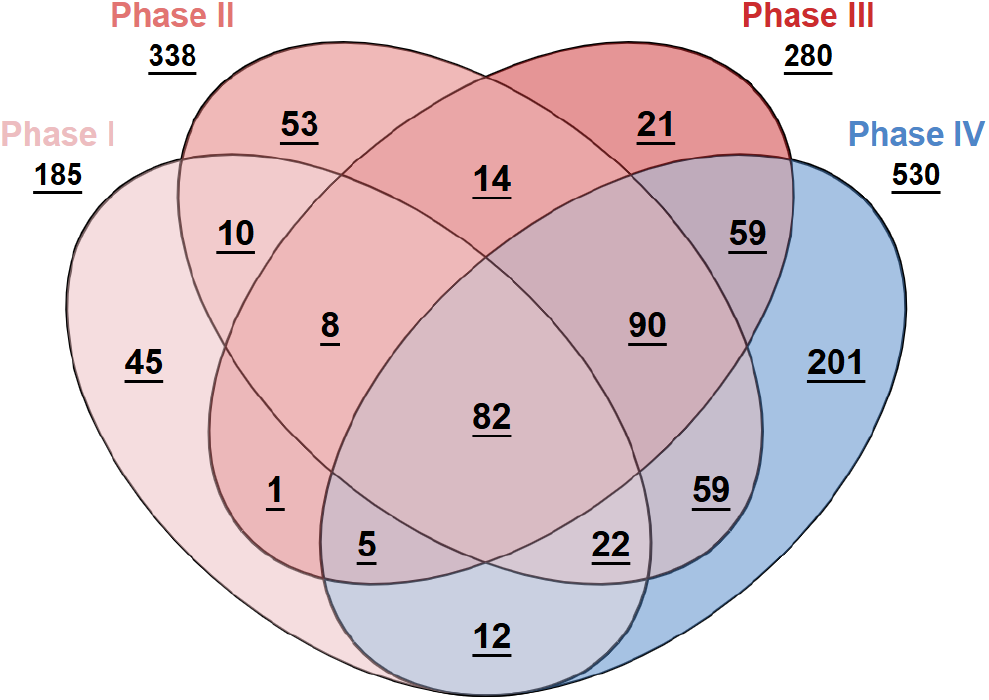
Agents/drugs by phase I-IV. Agents and drugs can be intersected by the four clinical phases via a Venn diagram. A single compound may appear in multiple phases. The same functionality is also offered for targets (not shown here).

This tool can pinpoint drugs that are already approved (in phase IV) and are also currently being repurposed for new indications in one or more of the clinical trial phases I-III. For example, terazosin is an approved antagonist of α_1A/1B/1D_–adrenoceptors for hyperplasia of prostate and essential hypertension. Additionally, terazosin is currently being tested in Phase I and Phase III clinical trials for treatment of Parkinson’s disease and diseases of the circulatory system, respectively. Furthermore, the tool can provide information about agents that are not yet approved but are broadly investigated in clinical trials. A point in case is atrasentan that is an endothelin A receptor antagonist which is not approved while being studied in clinical trials across all phases I-III for the treatment of several types of neoplasms and IgA nephropathy. Thus, this tool helps to find both drugs being repurposed as well as new agents that are awaiting their first approval.

### Targets

#### Target page

This page describes a drug or clinical trial target, which can be searched for by its gene or protein name. It offers a filterable tabulation of all the target’s compound–disease pairs preceded about information about the activity at different clinical phases (max phase and individual phases). The compound information includes name, molecule type and pharmacological modality. The disease information spans indication names, ICD11 codes, ATC codes and association scores (from Open Targets). For example, when searching for the GIP receptor, we can find it is targeted by eight different compounds, one approved drug (tirzepatide) and seven agents in clinical trials. Tirzepatide (Mounjaro) was approved in 2022 for obesity and type 2 diabetes mellitus, and it is being investigated in phase II for non-alcoholic steatohepatitis. Additionally, seven new agents are being studied in phase I-III for the same indications, these compounds are mostly proteins, except one small molecule and one antibody.

#### Targets by phase I-IV

This page intersects targets by any combination of clinical phases. As for compounds (above), clicking areas of a Venn diagram updates a table of targets, which are described by their gene and protein names, and classification (class, ligand type and receptor family sharing physiological ligand). Compounds are described by their name, molecule type and pharmacological modality. The disease indication information includes disease areas and diseases annotated using the ICD11 and ATC (drugs only) classifications. Thus, the starting point is targets in one or more clinical phases leading while further focusing is possible based on any combination of compound and disease information.

This tool can help select GPCRs, currently 83 receptors, that are being re-targeted i.e., have approved drugs and are again investigated in clinical trials with other agents and/or disease indications. For example, the chemokine receptor CXCR4, is targeted by three drugs and three agents, and has 22, 14, one and five indications presently, in phase I, II, III and IV, respectively. Conversely, the tool can also identify novel targets i.e., GPCRs are being targeted by agents in clinical trials but lack approved drugs. There are 30 such novel targets, including CXCR1 and CXCR2 that are currently being investigated in all of phases I, II and III. This signifies the substantial interest in these targets and their potential to become the targets of approved drugs. Furthermore, the Venn diagram can reveal which targets have spurred the latest interest for drug development. Specifically, 12 receptors have agents only in Phase I clinical trials making them the newest targets of clinical investigation. Altogether, this tool informs of the current status of specific GPCRs as drug targets pinpoints both those being re-targeted and entering clinical trials for the first time.

#### The drugged GPCRome

This page visualizes the clinical status of different GPCRs. The first tab shows a GPCRome wheel that covers all 398 human non-odorant GPCRs (Fig. 4a). The GPCRome wheel is a new plot that is only available in GPCRdb and is tailored for the mapping of diverse data types, including those uploaded by users^16^. Here, each ring contains one or two GPCR classes divided into receptor families (by physiological ligand) and then individual receptors – all of which are sorted alphabetically. This enables quick look up of a given GPCRs and spatial grouping of related receptors. The clinical status is depicted in a color-coded ring segment inside each receptor name. Beyond the phases I (light pink), II (light red), III (bright red) and IV (blue), this also includes untargeted receptors with (grey) or without (white) disease association as well as sensory GPCRs (light cyan). The threshold for disease associations is 0.5 and represents a maximum value across all diseases imported from the Open Targets database^20^. Altogether, the GPCRome wheel gives an overview of the level of clinical investigation across the complete non-odorant repertoire of GPCRs in human.

**Figure 4.**
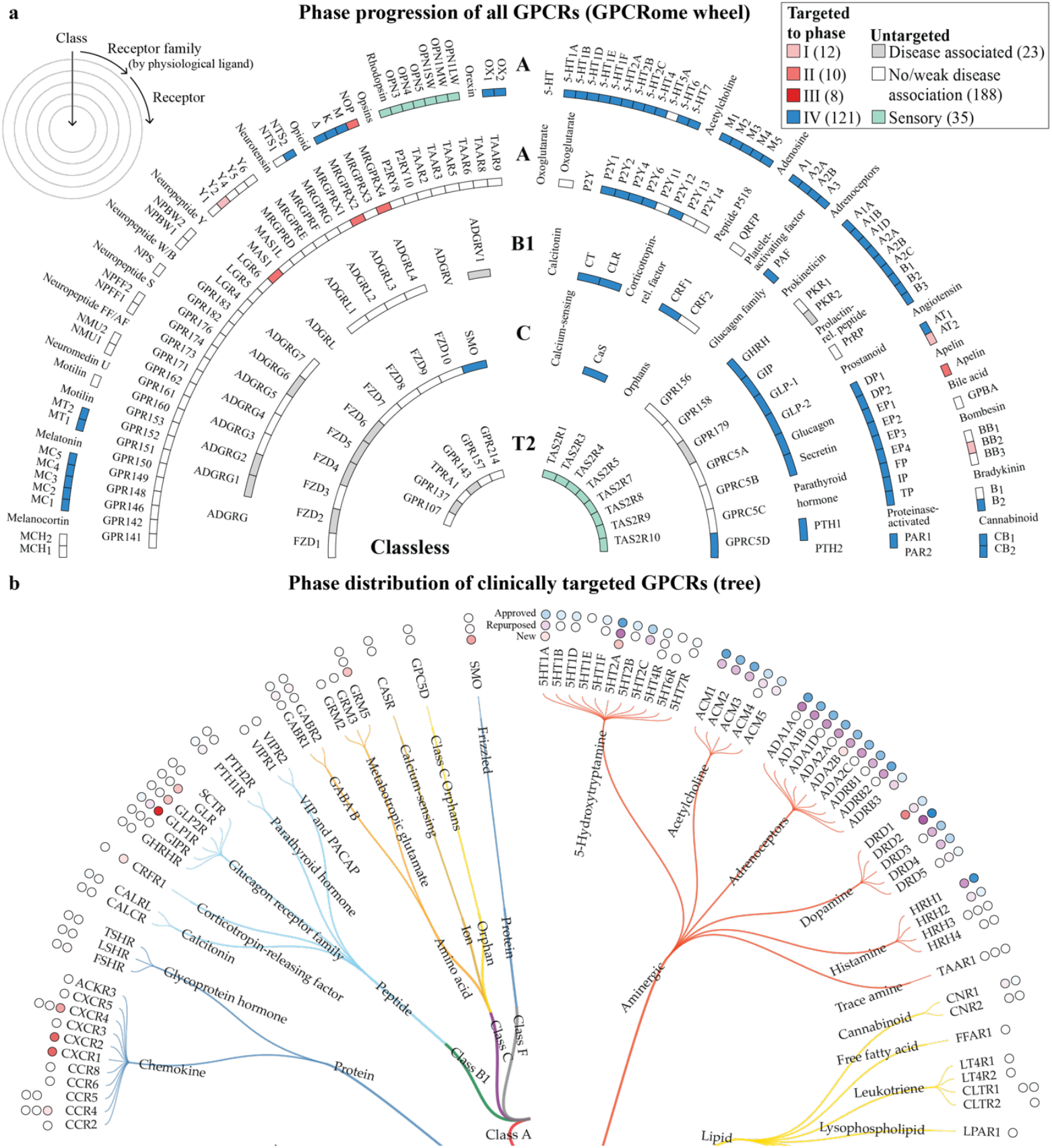
The drugged GPCRome. **a**, GPCRome wheel showing targeting status of all non-odorant GPCRs, which are grouped by class across concentric layers. Each receptor is color-coded according to the highest clinical phase of all compounds targeting it. Untargeted GPCRs are subdivided into disease associated or no/weak disease association whether they have or not at least one disease with association score (from OpenTargets) equal or higher than 0.5. Sensory receptors are colored in green. **b**, Tree showing the distribution across classes and receptor families of 151 GPCRs targeted by approved drugs (121) or agents in clinical trials (30). Each receptor is depicted with up to four external circles, representing the number of compounds in phase I – IV, from the innermost to the outermost circle, where data is available. The circles are color-coded based on two different gradients, one based on the number of agents in clinical trials (phase I – III) and another one based on approved drugs (phase IV).

The GPCRome wheel can be used to map drugged GPCRs (phase IV) across the different classes showing a distribution of 104 class A, 11 class B1, 5 class C and 1 class F receptors. Although the latter class, F only has a single drugged member, the smoothened receptor (SMO), it has four additional receptors that are associated with diseases and could expand the representation of this class in clinical investigation and eventually, clinical practice. Similarly, class B2, which lacks approved drugs altogether, has six receptors with associated diseases making it likely that this class too could enter clinical trials in the coming years. When focusing the analysis onto individual receptors that could soon become drugged, the chemokine receptors stand out with nine members currently in phases I-III. Interestingly, six additional class A GPCRs being targeted by agents in phase I-II are orphan receptors i.e., have unknown physiological ligands. This shows that target identification and drug discovery is possible even without knowledge of the endogenous activator mediating the physiological process.

The second and third tabs in the page *The drugged GPCRome* shows a tree that is focused on the subset of receptors that have been clinically targeted (Fig. 4b). Like in the GPCRome wheel, receptors are classified and sorted by class, receptor family and name, but in a single ring. The clinical status is shown in circles outside of the receptor names. The circles are filled with a gradient from light to dark by an increasing number of agents for the given receptor (the darkest shade is the max values in each ring of circles). In the first tree, three circles represent the number of novel agents in trials (red), approved drugs being repurposed in trials (purple) and all approved drugs (blue). The second tree instead has four circles each of which represent the number of compounds in a given phase I-IV. Hence, the first tree separates novel from repurposed compounds while the second tree describes the activity in a specific phase for a given GPCR target. This gives the trees a complementary utility relative each other and the GPCRome wheel when studying the progression of targets across the clinically targeted GPCRome.

One observation from the first tree is that aminergic receptors in class A have many approved drugs but relatively fewer agents in clinical trials, indicating a possible saturation of this receptor group^3^. For example, most members of the serotonin (5-Hydroxytryptamine) family have reached phase IV but seem to be of lower interest in ongoing drug development. However, they are still the most prominent receptor group in terms of drug repurposing, mostly in phase III trials, as seen in the second tree. This could be explained by the high number of drugs targeting them, especially in receptors such as dopamine D_2_ or 5-HT_2A_ receptors. Conversely, the Chemokine receptor family in class A is gaining interest in clinical investigations, having only three receptors targeted by marketed drugs while nine are currently in clinical trials. An intermediate example is the six receptors in the glucagon family in class B1, since these are all drugged and also increasingly targeted in clinical trials.

#### Target selection tool

The target selection tool (Fig. 5) serves to guide the section of targets for clinical investigation by efficiently navigating multi-modal data integrated from major drug, target and clinical trial database databases (Table 1) and manual annotation. The Target selection tool is structured as six tabs containing complementary data tables. The tables are inter-connected so that any filter or sorting applied based on one type of data is also carried on when switching view. The first tab, ‘Target characterization’ outlines the target novelty classification, number of literature articles and counts of inactive, active and total experimental structures (from Pharos^21^, PubMed <u>(www.ncbi.nlm.nih.gov/pubmed)</u> and PDB^22^, respectively). The second tab, ‘Compounds’ provides counts of the number of drugs, agents in current clinical trials and ligands with bioactivities (see Agent/drug page above). The next two tabs provide disease information for the clinically targeted and yet untapped receptors, respectively. The former tab contains disease areas and diseases (ICD11 classification), disease indications (ATC classification) and disease association scores of targets (from Open Targets). The latter tab contains only the target disease association scores, as untapped targets lack clinical agents. The last two tabs, describe tissue expression and cancer expression data for receptors (from the human protein atlas^23^). Together, these features make it possible to select specific targets of interest ranging those never previously explored in clinical trials to GPCRs with multiple drugs and remaining untapped disease indications.

**Table 1.** Data sources for drugs, agents, targets and disease indications and associations.

| Data sources | Drugs | Agents | Targets | Disease indications | Target disease associations |
| --- | --- | --- | --- | --- | --- |
| DrugBank<sup>26</sup> | X | X | X |  |  |
| Open Targets<sup>20</sup> | X | X | X | X | X |
| ClinicalTrials.gov | X | X |  | X |  |
| Manual annotation (web sites, press releases etc.) |  | X | X | X |  |

Regulatory agencies covered include the US Food and Drug Administration (FDA), European Medicine Agency (EMA), Japanese Pharmaceutical, Medical Devices Agency (PMDA) and more.

**Figure 5.**
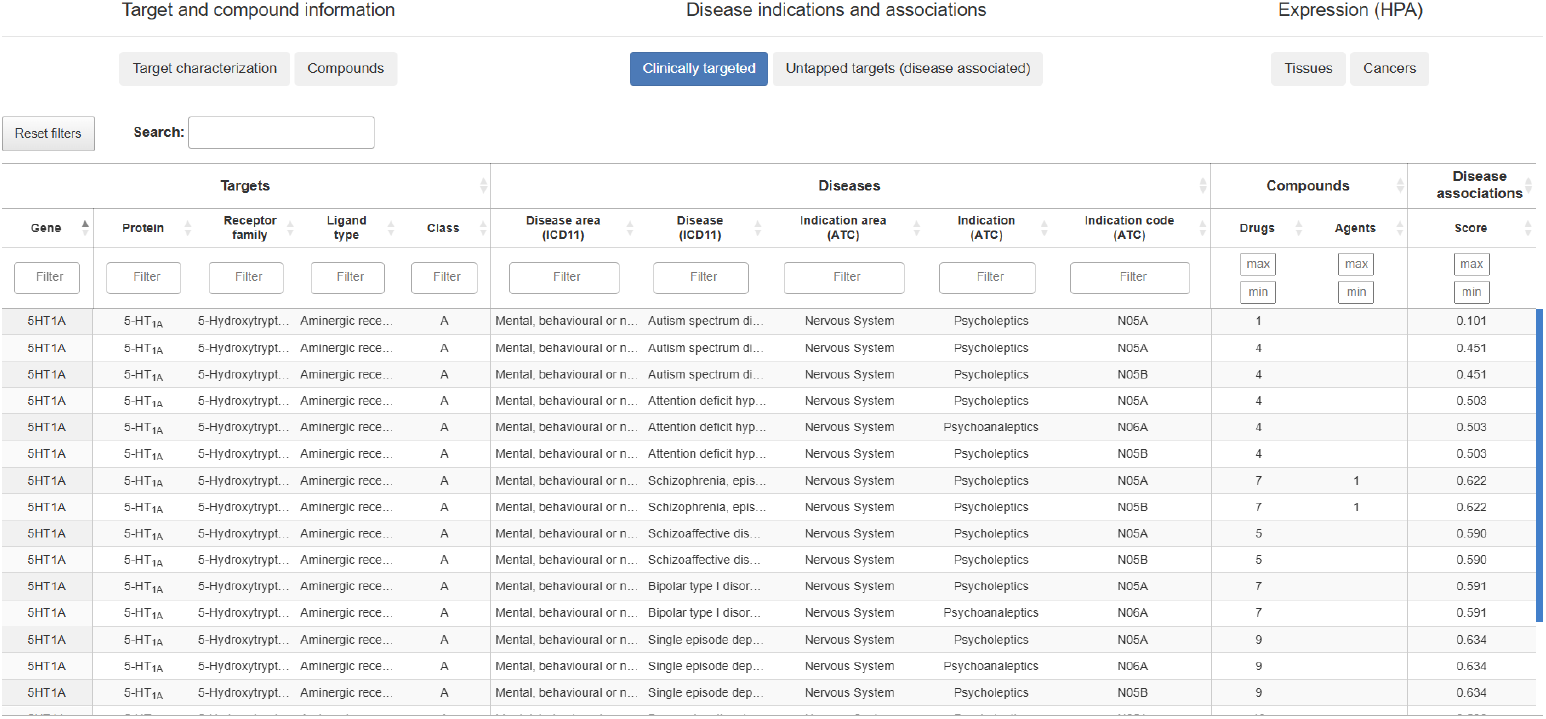
Target selection tool. The target selection tool offers a powerful yet swift means to prioritize GPCRs for future drug discovery based on disease indications, characterization, tissue expression and more. It supports studies looking for both novel targets as well as already drugged targets that may be targeted for new disease indications.

### Diseases

#### Disease page

The starting view of this page is a Sankey plot giving an overview of connections of the selected disease to all its associated targets agents/drugs along with their targets. Next, an interactive table shows the agents’/drugs’ progression across clinical phases (max, I, II, III and IV), molecule type (e.g., peptide or small molecule), pharmacological modality (e.g., agonist or antagonist), and target names and classification. Finally, another dynamic table provides multi-modal target information about targets. This spans disease associations (from Open Targets) based on genetic association studies and somatic mutations. Furthermore, it covers scores for known drugs (ChEMBL), literature publications (Europe PMC), RNA expression (expression atlas) and animal models (IMPC) (all explained in https://platform-docs.opentargets.org/evidence).

#### Disease overview

Our integration and annotation of disease data generated 23 major disease areas encompassing 683 disease indications classified using the ICD11 disease classification. To enable overview as well as more in-depth analysis of the large disease data across clinical trials and the market, we developed an interactive outline (Fig. 6a). This outline initially displays only the 23 disease areas, but each of these can be expanded multiple times to show a listing of child-terms and ultimately, specific diseases. The representation of diseases across current clinical trials and the market as shown in three color-coded circles visualizing the proportion of new agents, drugs being repurposed and all approved drugs for each disease and area (max value among encompassed diseases). Additionally, a second tab provides a multi-column listing showing a simultaneous view of all (275) disease categories one the second level (Fig. 6b). Together, these two tools offer a quick means to analyze relative frequencies and trends in disease indications of GPCR-targeted clinical compounds, and to visualize them in a single diagram figure publication or presentation.

**Figure 6.**
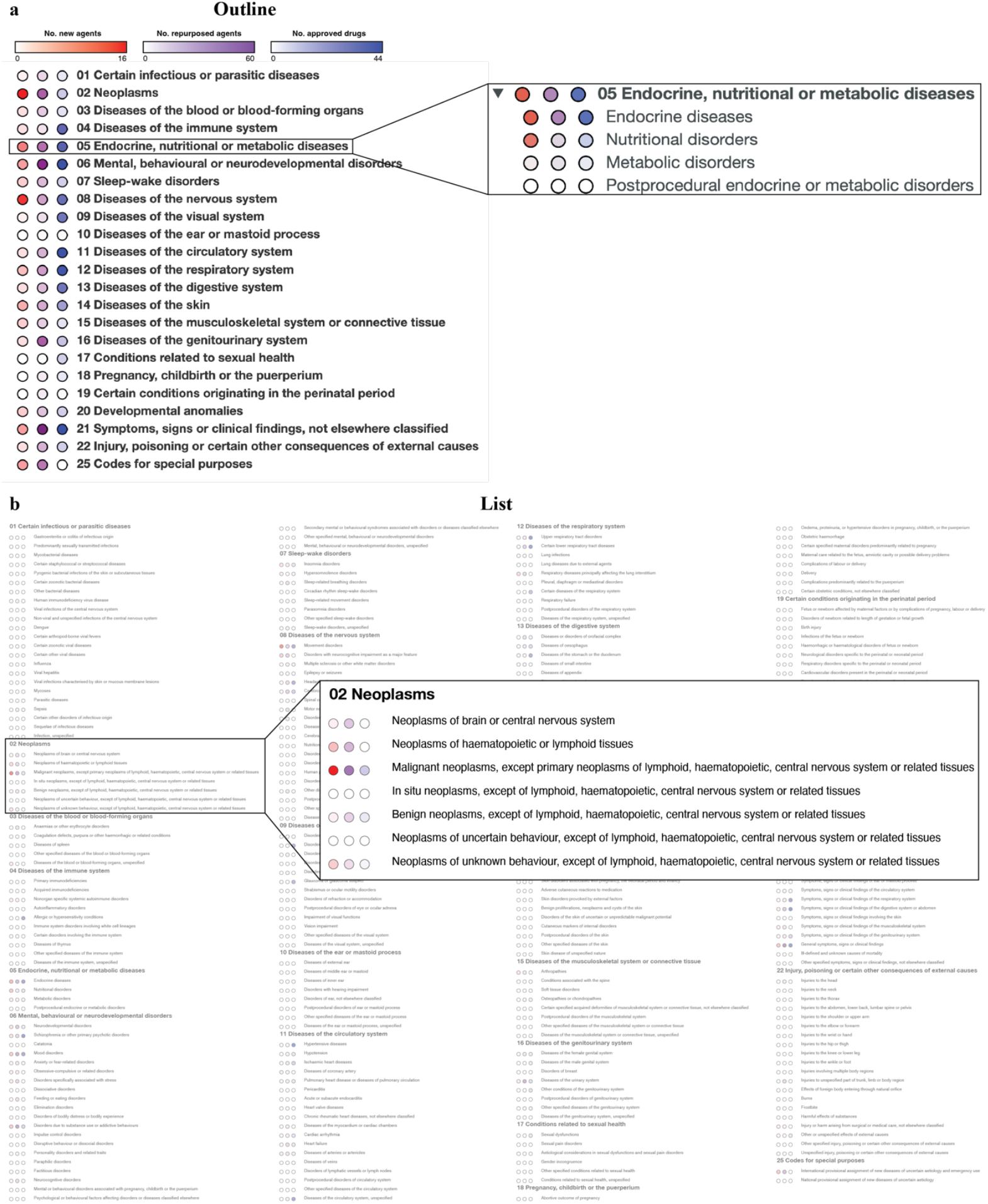
Disease overview. The *Disease overview* page showing the distribution of compounds across 683 diseases grouped into 23 areas – all of which are classified using the International Classification of Diseases (ICD11)^17^. The circle fill gradient is proportional to the number of new agents (red), drugs being repurposed (purple) and all approved drugs (blue). **a**, Outline of disease areas each of which can be expanded to show the set of subcategories and ultimately, diseases belonging to the area. **b**, List plot simultaneously showing all 275 disease categories one the second level of the disease classification.

## Discussion

Given their large abundance and extensive targeting, building a resource dedicated to drug discovery on GPCRs will benefit a large community globally. It offers a one-stop-shop for compounds (agents and drugs), targets and diseases (indications and associations) combining integration of several major databases with extensive manual annotation. Notably, the annotation of company web sites and press releases doubled the number of clinical trial agents from ClinicalTrials.gov linked to GPCR targets. Furthermore, the targets are enriched with data already curated in GPCRdb, such as structures, ligands^18^ and 36,000 ligand binding site mutations^24^. The resource enables data to be intersected in new ways. For example, the Venn diagrams of agents/drugs and targets can extract those being present in just a single phase or a specific combination of phases. Finally, also the visualization of data has been tailored to GPCRs. Most notably, the new GPCRome wheel^16^ presents a unified view of the progression of all non-odorant GPCRs across the yet untargeted and clinical phases. Furthermore, the two trees allow more in-depth analysis of the activity in each clinical phase, novel agents and repurposing. Taken together, the online resource focused on GPCRs complements major generalist databases such as DrugBank and Open Targets by offering unique integration, curation, analysis and visualization of a broad diversity of data. Its utility is demonstrated by our accompanying analysis revealing the latest trends in GPCR drug discovery^3^.

Selection of targets for drug discovery is typically a balance of numerous parameters, such as their characterization functionally and therapeutically, landscape of existing drugs or agents in trials, templates for structure-based design, expression across tissues and cancers etc. The Target selection tool interconnects all these types of data enabling users to step-by-step narrow down the GPCRs with the highest relevance and feasibility for the development of new medicines. Of note, this tool allows for a simultaneous selection across multiple targets and diseases (compound indications and experiment-based associations) not possible in other databases, including Open Targets. Furthermore, the versatility of the data and interface makes the tool applicable to both highly characterized targets that can be explored for new indications as well as novel targets never tested in clinical trials before. In future years, it will be beneficial to explore ways to expand the data further for example, by targeted annotation of literature and patents or a pre-competitive partnership with industry consortia. In terms of application of data-driven selection, it would be very valuable to explore drug repurposing, as was recently reported for DrugMechDB^25^.

## Methods

### Agent and drug data

We obtained the list of all drugs and agents in clinical trials targeting any non-sensory GPCR, alongside their mechanism of action and clinical indications from the public databases Open Targets^20^ and DrugBank^26^. We obtained the set of all drugs and agents targeting GPCRs from OpenTargets through programmatic access by iterating over each receptor. We downloaded the whole DrugBank dataset and only included those compounds with a known pharmacological action of a GPCR, or compounds without known pharmacological action but at least 50% or higher proportion of the targets being GPCRs. Both datasets were filtered out to remove compounds which have been withdrawn from the market or are nutraceutical or illicit. To obtain a unique set of drugs and agents, we combined entries from both resources by the ChEMBL identifier. In addition, to obtain a non-redundant set of molecules without duplicates of salt forms and other child molecules, we merged molecules under the same parent compound ID from PubChem^27^.

To obtain the most up-to-date set of agents in clinical trials, we only included agents from ongoing clinical trials, here defined as updated in the last three years and not failed or terminated. The primary resource for clinical trials is ClinicalTrials.gov, which currently consolidates over 500,000 trials worldwide. To ensure an up-to-date dataset (October 2024), we removed all agents that had been discontinued or removed from company websites. In addition, we included over 100 agents currently in clinical trials by manually annotating company websites and press releases. These compounds had at least one trial registered in ClinicalTrials.gov, but were not found neither in DrugBank nor in Open Targets.

### Disease data

We fetched Anatomical Therapeutic Chemical (ATC) codes from DrugBank. ATC codes describe organ/system of action, mechanism and/or chemical scaffold. To achieve another classification focusing on diseases and applicable not only to drugs but also agents in clinical trials, we used the International Classification of Diseases (ICD11)^17^. The ICD11 codes were obtained by using a coding tool^17^ to translate indications from OpenTargets^28^, which uses the EMBL-EBI Experimental Factor Ontology (EFO), the Human Phenotype Ontology (HPO) and the Mondo Disease Ontology (Mondo). Furthermore, these disease indications were manually verified for each compound-indication pair retrieved from ClinicalTrials.gov. To focus on drug development, we excluded the areas “Supplementary Chapter Traditional Medicine Conditions”, “Supplementary section for functioning assessment” and “Extension Codes”, as they do not contain relevant disease information. Additionally, the areas “External causes of morbidity and mortality” and “Factors influencing health status or contact with health services” were not present in our dataset and none or their diseases are being investigated or approved by GPCRs agents or drugs. Hence, our dataset covers 683 diseases indications across 23 areas. Target disease association scores of all non-sensory GPCRs were compiled from Open Targets through programmatic access^20^. The disease-target association score is a normalized weighted harmonic sum of seven evidence data sources: genetic associations, somatic mutations, known drug, affected pathway, literature, RNA expression and animal models.

### Additional target selection tool data

In addition to the abovementioned data, we imported data of particular relevance to the target selection tool. The target novelty score, alongside with the publication count and the target development level based on the Illuminating the Druggable Genome (IDG) were obtained from Pharos through programmatic access^21^. The target novelty score is computed based on the relative abundance of the target associated publication. The IDG target levels are Tdark, Tbio, Tchem and Tclin from less to more characterized and advanced targets in terms of ligands and drugs targeting them^29^. Additionally, we included tissue and cancer expression data from the Human Protein Atlas (HPA)^23^. The tissue expression data is based on transcriptomics data from HPA and GTEx^30^, which was calculated as the maximum normalized expression value for each gene and tissue among the two data sources. For tissues containing sub-tissues (brain, lymphoid and intestine), the maximum of all sub-tissues was used as the tissue type value. Cancer data is based on staining profiles through immunohistochemistry using tissue micro arrays in 20 human tumor tissues. For each tumor tissue, four different staining levels were defined (high, medium, low and not detected). For each GPCR-cancer tissue we obtained the max staining level identified. Both tissue and expression data were based on The Human Protein Atlas version 24.0 and Ensembl version 109.

### Programming framework

GPCRdb builds on the Django web framework version 2.2^31^ and uses PostgreSQL to manage and store data in the backend database. Connections between the PostgreSQL database and the frontend tables are managed via dedicated Python queries and classes, whereas graphical representations such as the Venn diagrams and the GPCRome wheel have been implemented using the JavaScript D3.js library.

## Data availability

GPCRdb is available at https://gpcrdb.org and can also be accessed via a RESTful API, which complies with the OpenAPI specification using Swagger (code examples are available at https://docs.gpcrdb.org/web_services.html). The underlying data and a virtual machine configuration are all available in the repositories at https://github.com/protwis.

## Code availability

The source code is available in the repositories at https://github.com/protwis.

## Acknowledgements

This work was supported by the Lundbeck Foundation [R383-2022-306], and the Novo Nordisk Foundation [NNF23OC0082561] to D.E.G.

## Author contributions

J.C, J.S.L. and D.E.G. wrote the main manuscript text. D.E.G. prepared Figure 1 and Table 1. B.X. prepared Figure 2. S.N.A. prepared Figures 3 and 5. J.S.L. curated the data and prepared Figures 4 and 6. All authors commented on the manuscript. J.C., S.N.A., B.X. and G.P.S. developed the online resource. D.E.G. conceptualized the study, raised the funding and supervised and administered the work.

